# Modeling the critical MCOR-causing deletion in mouse unveils aberrant *Sox21* expression in developing and adult iris and ciliary body, and implicates *Tgfβ2* in MCOR-associated glaucoma and myopia

**DOI:** 10.1101/2023.12.11.571097

**Authors:** Clémentine Angée, Elisa Erjavec, Djihad Hadjadj, Bruno Passet, Pierre David, Corinne Kostic, Emmanuel Dodé, Xavier Zanlonghi, Nicolas Cagnard, Brigitte Nedelec, Sylvain V. Crippa, Christine Bole-Feysot, Mohammed Zarhrate, Sophie Creuzet, Johan Castille, Jean-Luc Vilotte, Patrick Calvas, Julie Plaisancié, Nicolas Chassaing, Josseline Kaplan, Jean-Michel Rozet, Lucas Fares Taie

**Author notes:** Contributed equally (Clémentine Angée, Elisa Erjavec).

## Abstract

Congenital microcoria (MCOR) is a rare hereditary developmental defect of the iris dilator muscle, frequently associated with high axial myopia and high intraocular pressure (IOP) glaucoma. The condition is caused by submicroscopic rearrangements of chromosome 13q32.1. However, the mechanisms underlying the failure of iris development and the origin of associated features remain elusive. Here, we present a 3D architecture model of the 13q32.1 region, demonstrating that MCOR-related deletions consistently disrupt the boundary between two Topologically Associating Domains (TADs). Deleting the critical MCOR-causing region in mice reveals ectopic *Sox21* expression precisely aligning with *Dct*, each located in one of the two neighbor TADs. This observation is consistent with the TADs’ boundary alteration and adoption of *Dct* regulatory elements by the *Sox21* promoter. Additionally, we identify *Tgfb2* as a target gene of SOX21 and show TGFB2 accumulation in the aqueous humor of a MCOR-affected subject. Accumulation of TGFB2 is recognized for its role in glaucoma and potential impact on axial myopia. Our results highlight the importance of SOX21-TGFB2 signaling in iris development and control of eye growth and IOP. Insights from MCOR studies may provide therapeutic avenues for this condition but also for glaucoma and high myopia conditions, affecting millions of people.

## INTRODUCTION

The iris is a remarkable ocular structure, serving a dual purpose by finely tuning the amount of light that reaches the retina and regulating intraocular pressure (IOP). It comprises a stroma, a double epithelium layer and two dynamic muscles, the dilator and the sphincter, that work together to adjust the size of the pupil and hence the light intensity for optimal vision. The iris is rooted to both the corneal-scleral junction and the ciliary body (CB), whose stroma and epithelial layers are continuous with those of the iris. This structural arrangement creates an open space known as the irido-corneal angle where the aqueous humor (AH) produced by the CB to nourish ocular tissues is drained out, playing a vital role in maintaining IOP.^1^ An increase in IOP is a prominent risk factor for optic nerve damage and glaucoma (GLC).^2^

Congenital microcoria (MCOR, MIM#156600; reviewed by Angee *et al.*^3^) is an extremely rare autosomal dominant condition affecting both of these functions. It is characterized by the partial or complete absence of the iris dilator muscle, which normally originates from the anterior part of the epithelial layer (AEL) and extends along its stroma. The sphincter muscle near the pupil’s edge retains full functionality. The absence of dilator muscle fibers is evident in the presence of pinhole-sized pupils (< 2mm) that exhibit minimal or no dilation, even when mydriatic drugs are used. This also leads to the iris thinning and atrophy of the stroma, particularly in the dilator muscle region, which permits light to pass through, inducing iris transillumination.^3^ In the vast majority of the cases, this abnormal development of the iris does not affect the chamber angle that is wide-open although prominent iris processes and a higher insertion of the iris root in the angle have been typically described.^3^ Alongside dilation problems, individuals with MCOR are at a substantially increased risk of developing high axial myopia, affecting 70% of cases.^3^ Among those with high myopia, half also experiences juvenile-onset high IOP GLC, an unusually high prevalence compared to the general population, which stands at 34% of all MCOR cases.^3^ The underlying causes of these associated features is highly debated. It was suggested that the tendency to close the eyelids to reduce the glare caused by iris transillumination in MCOR could lead to form-deprivation myopia. However, the absence of a clear link between myopia and oculocutaneous albinism, which shares similar symptoms of iris transillumination and light sensitivity, challenges this notion.^3^ In relation to GLC, high axial myopia (a known contributor to the condition),^4,5^ and the anatomical features of microcoric chamber angles, which may affect AH flow, may play a contributory role but are not solely sufficient. This is evident from the fact that although all individuals with GLC exhibit these anomalies, only about a third of those with these anatomical variations actually develop GLC.^3^

The disease was ascribed to submicroscopic rearrangement of chromosome 13q32.1, particularly deletions and a reciprocal duplication, suggesting that the disease is linked to an alteration of the regulatory landscape of the region.^6–8^

Here, we provide a 3D architecture model of the 13q32.1 region that suggests that MCOR-related deletions consistently alter the boundary between two interacting Topologically Associating Domains (TADs). Deleting the critical 35 Kb MCOR-causing region in mice leads to aberrant *Sox21* (MIM*604974) expression in developing and adult iris and CB, aligning precisely with *Dct* (MIM*191275), that is located in the adjacent TAD. This suggests ectopic *Sox21* expression may stem from *Dct* regulatory element adoption, supporting the alteration of TAD boundaries.

Additionally, we identify the transforming growth factor, beta (TGFB)-2 encoding gene (*Tgfβ2*; MIM*190220) as a SOX21 target gene and TGFB2 accumulation in the AH of a MCOR-affected subject. This accumulation, known for its role in GLC^9–12^ and potential impact on axial myopia,^13,14^ underscores the significance of SOX21-TGFB2 signaling in these ocular anomalies and iris development. Insights from MCOR studies may offer therapeutic avenues for individuals with this condition, as well as those with primary open-angle glaucoma (POAG) and high myopia, affecting millions.

## RESULTS

### Deletions causing MCOR are predicted to modify the boundary between two adjacent topologically associated domains on chromosome 13q32.1

The MCOR locus, defined by its most distant deletion boundaries, spans 99.8 Kb pairs on chromosome 13q32.1 (GCRCh37/hg19_chr13: 95,209,609 – 95,309,380).^3^ According to Hi-C sequencing data available on the 3D Genome Browser, this locus was originally situated within a one megabase (Mb) TAD, encompassing the genomic region extending from *DCT* (MIM*191275) to *UGGT2* (MIM*605898). Utilizing higher-resolution Hi-C sequencing data obtained from mouse neural progenitor cells, we refined the structural organization of the 13q32.1 region through the detection of two interacting TADs of 1.2 (TAD1) and 1.5 Mb (TAD2), respectively (**Figure 1A and 1B**). TAD2 that comprises the original 1 Mb TAD can be broken down into three interacting sub-TADs of 0.2 (TAD2.1), 0.5 (TAD2.2), and 0.2 (TAD2.3) Mb, respectively. Remarkably, all documented deletions associated with MCOR (reviewed by Angée *et al.*^3^), exhibit modifications at the interface between TAD2.1 and TAD2.2, along with alterations within the regions of both TADs that engage in mutual interactions (**Figure 1A and 1B**). This underscores the significance of the boundary between TAD2.1 and TAD2.2 and emphasizes the intricate interplay within these topologically associating domains in the context of MCOR. None of the reported deletions have an impact on TAD1 or subTAD2.3 (**Figure 1A and 1B**).

**Figure 1:**
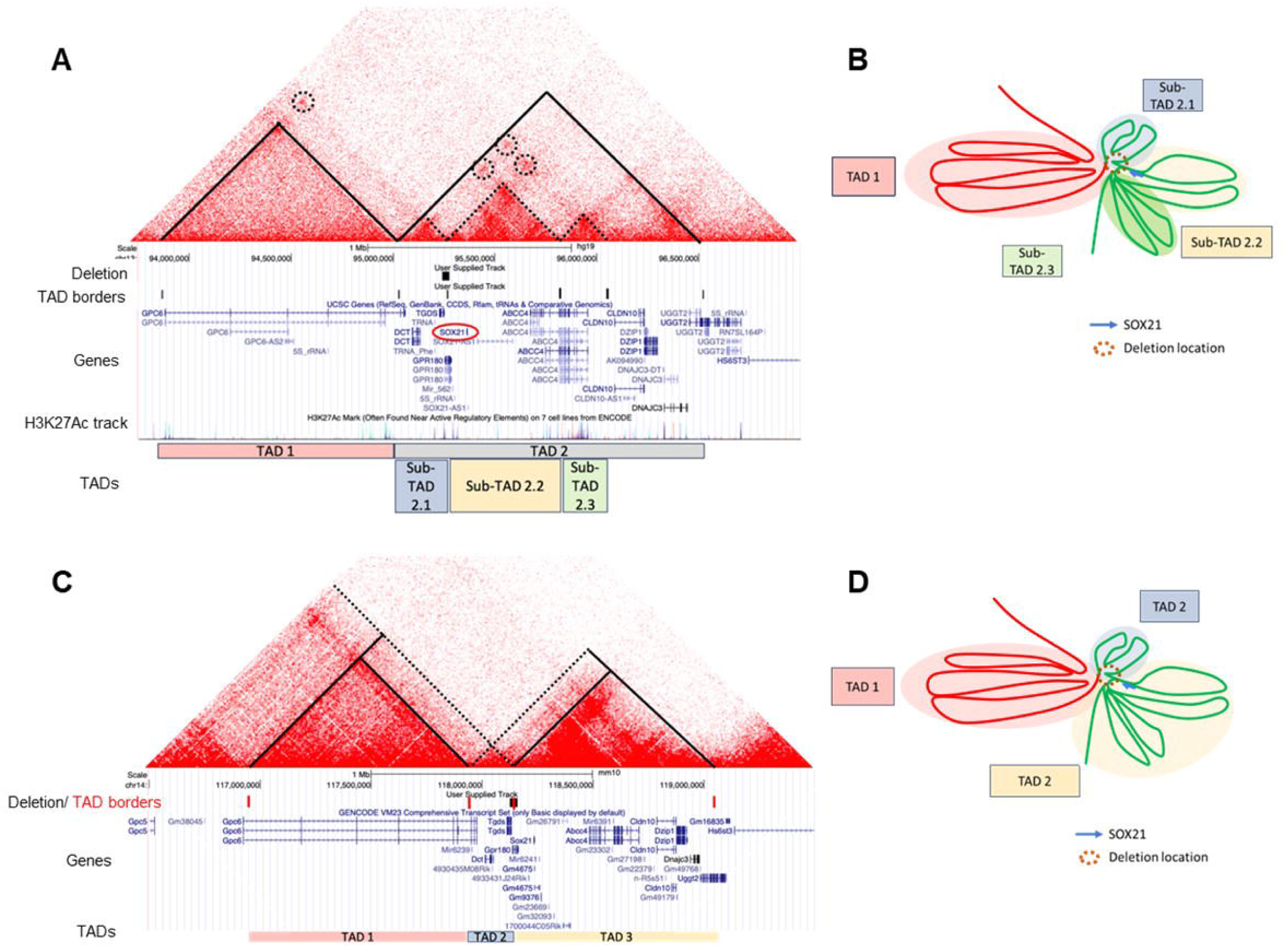
Organization of the MCOR Region in the human and syntenic mouse genomes. A) The 13q32.1 MCOR locus in humans (GCRCh37/hg19) is depicted. Based on HiC-seq data from human neural progenitor cells, the region consists of two topologically-associated domains (TADs) spanning 1.2 Megabases (Mb) and 1.5 Mb (in pink and grey, respectively). These TADs are demarcated by black straight lines on the interaction frequency map, with TAD borders indicated in the second track under the interaction map. TAD 2 further reveals at least three discernible sub-TADs: Sub-TAD 2.1 (in blue) covering 0.2 Mb, Sub-TAD 2.2 (in yellow) spanning 0.5 Mb, and Sub-TAD 2.3 (in green) covering 0.2 Mb. Sox21 is located within Sub-TAD 2.2, and the deletion affects the boundary between Sub-TAD 2.1 and Sub-TAD 2.2. Circled dashed lines highlight direct and robust interactions between the deleted region and other regions in various TADs and sub-TADs. **B)** The schematic representation illustrates the chromatin organization at the 13q32.1 locus. TAD 1 and the three Sub-TADs are emphasized in their respective colors. A dashed brown circle line denotes the location of the MCOR deletion, while the blue arrow highlights the Sox21 gene’s position in Sub-TAD 2.2. **C)** and **D)** depict the predicted 3D structure of the mouse syntenic region on chromosome 14. This structure closely mirrors the human configuration, demonstrating a high degree of similarity between the two species.

### The human MCOR locus at 13q32.1 aligns with a broad syntenic region on mouse 14qE4, covering TADs 1 and 2

Through a comparison of human and mouse genomic sequences, we demonstrate that TADs 1 and 2 reside within a large block of synteny shared by human chromosome 13q32.1 and mouse chromosome 14qE4, with minimal changes in gene composition and organizational structure occurring outside the subTADs 2.1 and 2.2 (**Figure 1C and 1D**). Based on the observed synteny, we generated C57BL/6 transgenic mice by using the CRISPR-Cas9 methodology to specifically delete the critical MCOR sequence (B6.cΔ MCOR mouse model where cΔ refers to the deletion of the GRCm38/mm10_chr14:118,125,917 – 118,160,688 genomic region). This approach was undertaken to investigate its regulatory architecture and to gain insights into its role in iris development, as well as its potential connections to GLC and high myopia.

### Homozygosity for the critical MCOR causing deletion is lethal in C57BL/6 mice

Mating of heterozygous B6.cΔ MCOR/+ failed to produce homozygous B6.cΔ MCOR/cΔ MCOR animals. Collected of embryos from 8.5 to 10.5 embryonic days (E) showed that homozygous animals ceased developing around E8.5 and regressed by E9.5 (not shown). Further analysis of E9 embryos revealed that homozygous mutants displayed noticeable abnormalities, including smaller size, varying degrees of cardiac hypertrophy, and one individual exhibited pericardial effusion consistent with compromised cardiac function. Additional anomalies included hypoplastic optic and otic vesicles, enlargements in specific facial structures, and irregularities along the body axis (**Figure S1A**).

For in-depth analysis of these abnormalities in homozygous embryos, serial histological sections were conducted (**Figure S1B**). Mutant embryos exhibited pyknotic nuclei in key regions, including the neural tube, hindbrain, and neural crest-derived tissues such as branchial arches and spinal primordia. Massive pyknosis was observed in these neural crest-derived tissues. Additionally, scattered pyknotic cells were found in mesenchyme, neural tube, and endodermal regions throughout the embryos (**Figure S1B)**. Although the mutant embryos displayed a well-regionalized and differentiated heart, the presence of red blood cells was noticed in the pericardium, potentially indicating myocardium issues. While the myocardium in mutants was thinner compared to controls, their developmental delay complicated conclusive assessments of abnormal differentiation (**Figure S1B)**. Of note, placenta sections showed no apparent differences between mutants and controls (not shown), suggesting that observed cardiovascular abnormalities are not linked to placental defects. Together these observations, in particular the pattern of pyknosis in mutant embryos, support defective neural crest migration and differentiation.^15^

### Heterozygosity for the critical MCOR causing deletion causes reduced pupil size

In contrast to homozygous animals, heterozygous littermates (B6.cΔ MCOR/+; referred to thereafter as the MCOR-mouse model) developed normally (**Figure S1A**) and B6.cΔ MCOR/+ irises were not transilluminable. Pupil response to light is not affected (**Figure 2**) however a moderate but statistically significant (p<0.01) reduction in base-line pupil size compared to B6.WT (Wild-Type) littermates was observed (**Figure 2**).

**Figure 2:**
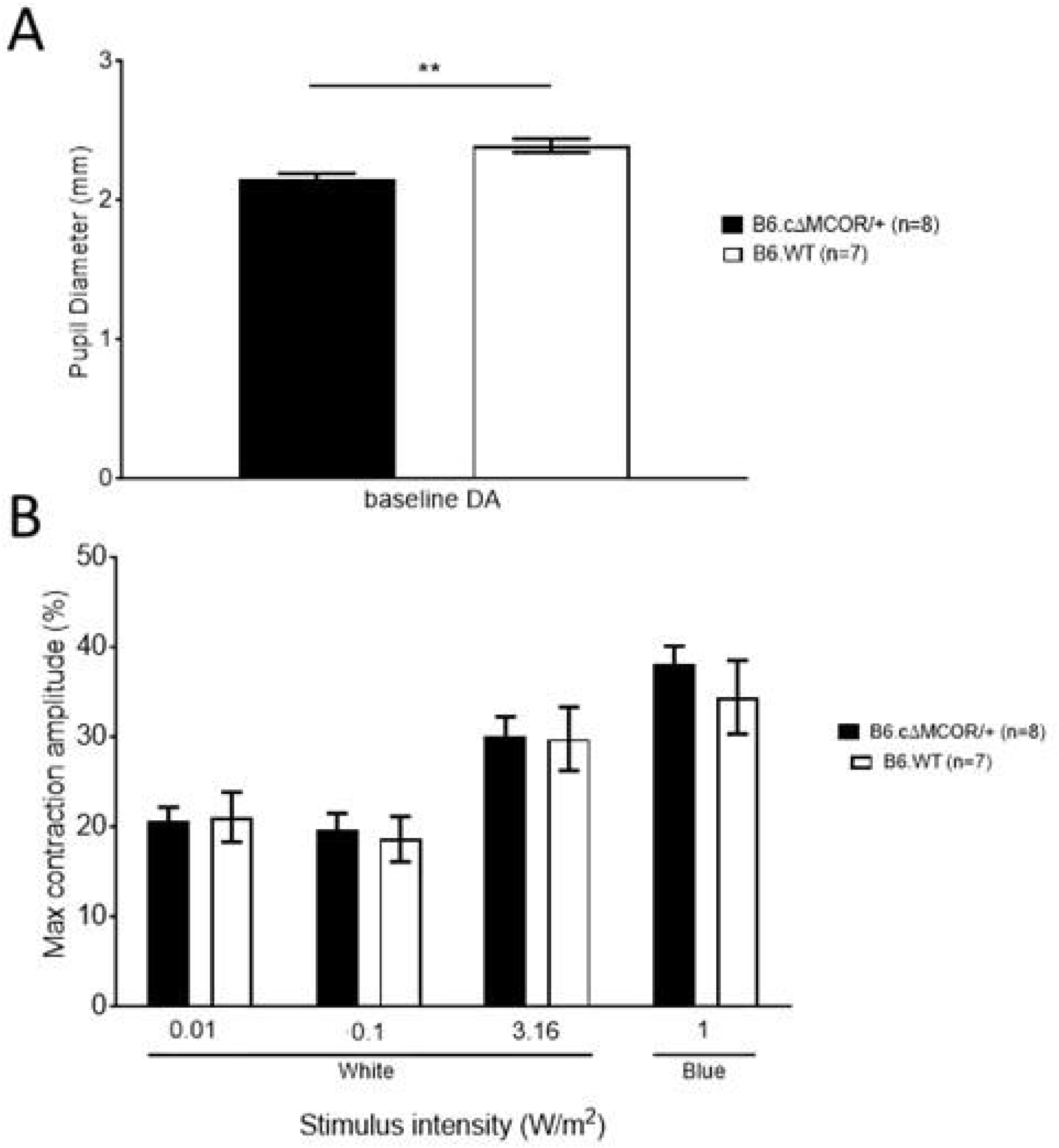
**Pupillary response of c**Δ**MCOR Mice. A)** A pupillometer was employed to assess pupil diameter in B6.cΔ MCOR/+ mice, revealing a moderate but statistically significant reduction in baseline pupil size compared to B6.WT animals (**: p<0.01, n = 8 animals per group). **B)** The quantification of the relative pupil size in response to white (0.01, 0.1 and 3. 16 W/m2) and blue (1 W/m2) light stimuli, revealed that the maximal contraction amplitude of B6.cΔ MCOR/+ mice in not significantly different from B6.WT mice in these conditions.

### *Sox21* in TAD2.2 and its product are ectopically expressed in iris tissue of the MCOR mouse model

To examine the impact of the critical MCOR-causing deletion on gene expression in iris tissue, we performed transcriptome sequencing on newborn B6.cΔ MCOR/+ and B6.WT irises. Consistent with heterozygosity for the deletion which alters *Tgds* (MIM*616146) and *Gpr180* (MIM*607787), the mRNA levels of these two genes in B6.cΔ MCOR/+ animals were approximately half of those found in B6.WT specimens (0.49 and 0.60, respectively; p < 10e-4).

Further analysis of the genes localized in the three predicted interacting subTAD within TAD2 revealed *Sox21* expression in irises from B6.cΔ MCOR/+, while no expression was observed in B6.WT littermates (p < 10e-9). This observation strongly suggests that the critical MCOR-causing deletion induces ectopic expression of *Sox21* (MIM*604974), a gene encoding a transcription factor belonging to the SRY-related HMG-box family^16^ located in subTAD2.2, approximately 50 kb downstream of the 3’MCOR locus boundary (**Figure 1**). The expression of the other genes located in the interacting TADs displayed comparable expression in mutant and WT irises (not shown).

RT-qPCR and Western blot analyses were conducted on iris RNA and protein extracts from adult mice using specific primers for *Sox21* and a highly specific SOX21 antibody,^17^ respectively. These analyses confirmed the sustained ectopic expression of *Sox21* in adulthood (**Figure 3A**) and disclosed the presence of a protein product in B6.cΔ MCOR/+ mice, conspicuously absent in B6.WT littermates (**Figure 3B and C**).

**Figure 3:**
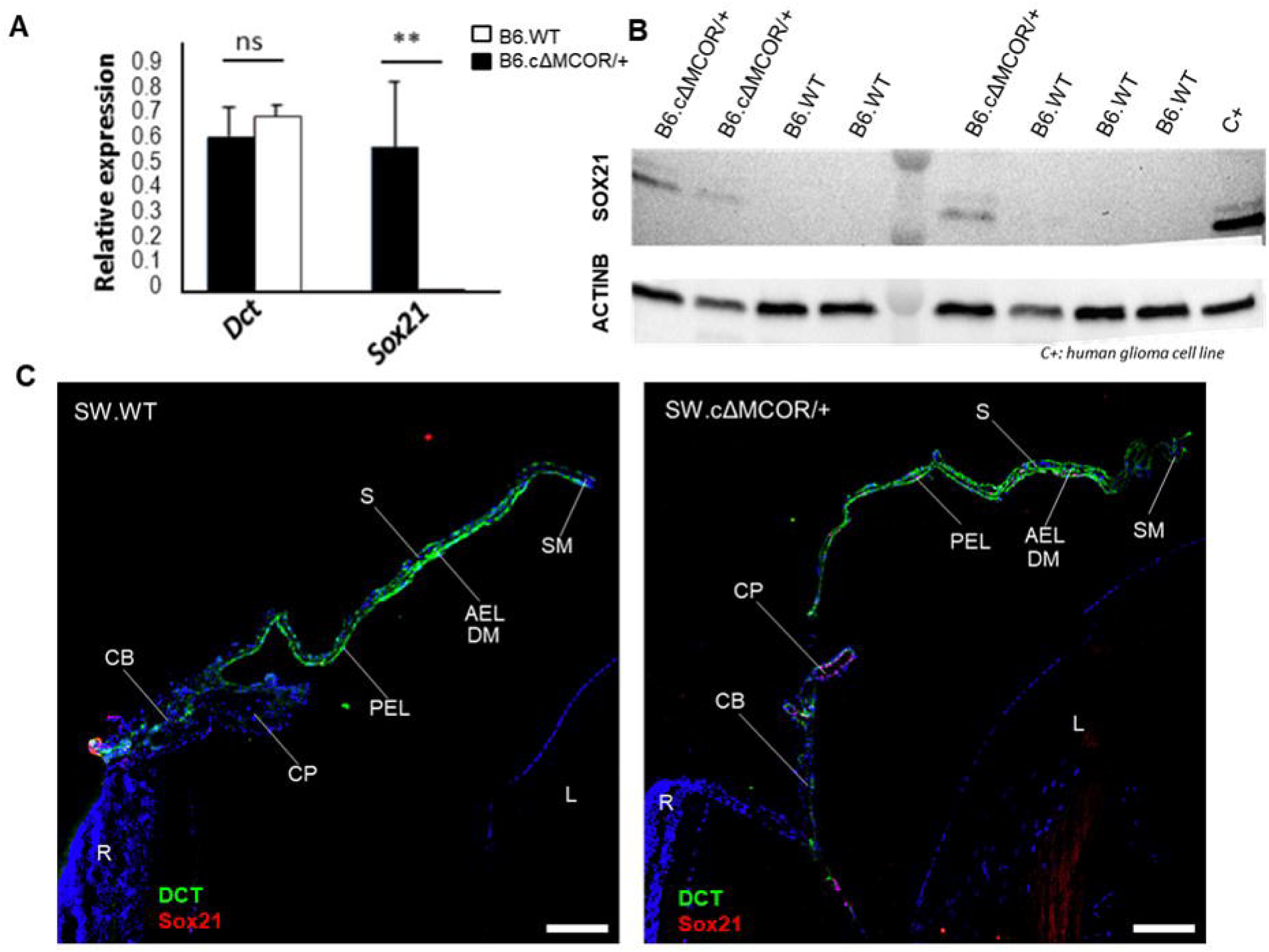
Expression Level Analysis at the 1Mb-TAD, RTqPCR, WB, and HiC Analysis of *Sox21* and its protein product in cΔ**MCOR and WT Animals. A**) RTqPCR analysis of *Dct* and *Sox21* abundance in iris/CB RNA extracts from newborn B6.cΔMCOR/+ and B6.WT mice (n = 5, each group). **B**) Western blot analysis of iris protein extracts reveals the presence of SOX21 in B6.cΔMCOR/+ but not in B6.WT counterparts. ACTB serves as the loading control, and C+ denotes glial cells expressing endogenous SOX21. **C**) Immunohistochemistry (IHC) analysis of iris and CB in newborn B6.cΔMCOR and B6.WT mice, illustrating the detection of SOX21 and DCT in their PELs. **AEL** – anterior epithelium layer, **CB**: ciliary body, **CP**: ciliary process, **DM:** dilator muscle, **L:** lens, **PEL:** posterior epithelium layer, **R:** retina, **S:** iris stroma, **SM:** sphincter muscle. (Scale bars: 100 µm).

### Sox21 is specifically detected in cells expressing DCT in the iris and ciliary body of the MCOR-mouse model

Immunohistochemistry (IHC) analysis of the iris in C57BL/6 animals is challenging due to the high pigmentation of the tissue. To circumvent potential alterations to iris integrity caused by depigmentation protocols, we opted to generate albino cΔ MCOR/+ and WT lines by mating B6.cΔ MCOR/+ animals with tyrosinase (TYR, MIM*606933)-negative Swiss albino (CFW) mice. RT-qPCR analysis of *Sox21* expression in irises of 2-month-old animals from the resulting lines, referred to as SW.cΔ MCOR/+ and SW.WT, revealed continued ectopic expression, despite the inactivation of tyrosinase (data not shown). We performed IHC analysis on iris and CB sections from adult animals using antibodies specific to SOX21 and DCT antibodies, respectively. *DCT*, also recognized as *TYRP2*, encodes the dopachrome tautomerase (EC 5.3.2.3). It operates as a melanogenic enzyme located just downstream of tyrosinase and serves as a distinctive marker for melanocyte lineages, ^18–19^ encompassing the pigmented epithelium layers of the iris and CB, respectively the iris posterior and CB anterior layers. *Dct* is positioned in TAD2.1 and a mere 75 Kb upstream of the 3’-boundary of the MCOR locus (**Figure 1A and B**). IHC exhibited robust staining of DCT in both the iris posterior and CB anterior epithelia in both SW.cΔ MCOR/+ and SW.WT animals (**Figure 3C**). This observation indicates that neither the deletion of the critical MCOR region nor the inactivation of tyrosinase has any discernible effect on the expression of *Dct* in albino mice. Interestingly, in SW.cΔ MCOR/+ animals, *Sox21* expression was limited to *Dct*-expressing epithelium layers of the iris and CB (**Figure 3C**). At this age, we observed no SOX21 staining in the AEL of the iris, which is the region from which the dilator muscle forms during the embryonic life (**Figures 3C**). Consistent with the ectopic expression of *Sox21* in mutant mice, SOX21 was undetectable in the iris and CB of SW.WT littermates (**Figure 3C**).

Combining RNAscope™ *In Situ* Hybridization (ISH) of *Sox21* mRNA and immune staining of DCT, we investigated their spatio-temporal pattern of expression from 10 embryonic days (E) to birth (postnatal day 0, P0) in SW.WT and SW.cΔ MCOR/+ eyes (**Figure 4**). In WT eyes, *Sox21* mRNA was detected in the proximal part of the optic vesicle (OV) at E10 and in the dorsal neuroectoderm at E10.5 and E11. However, from E11.5 onwards, although the DCT protein became evident, *Sox21* mRNA was no longer detectable, except in the surface ectoderm forming the eyelid and hair follicles, beginning around E15 (**Figure S2**). In mutant eyes, *Sox21* mRNA was initially detectable in the proximal part of the OV at E10 and in the dorsal neuroectoderm at E10.5 and E11, similar to its expression in WT littermates. However, from E10.5 onwards, *Sox21* mRNA was observed in the outer layer of the optic cup and later in the pigmented structures derived from this layer (**Figure 4A**). These pigmented structures in the developing eye include the retinal pigmented epithelium (RPE), as well as the AELs of the iris (reversed in adults) and CB (**Figure 4B**). They consistently expressed the DCT protein, which became readily detectable from E11.5 onward (**Figure 4A and B**).

**Figure 4.**
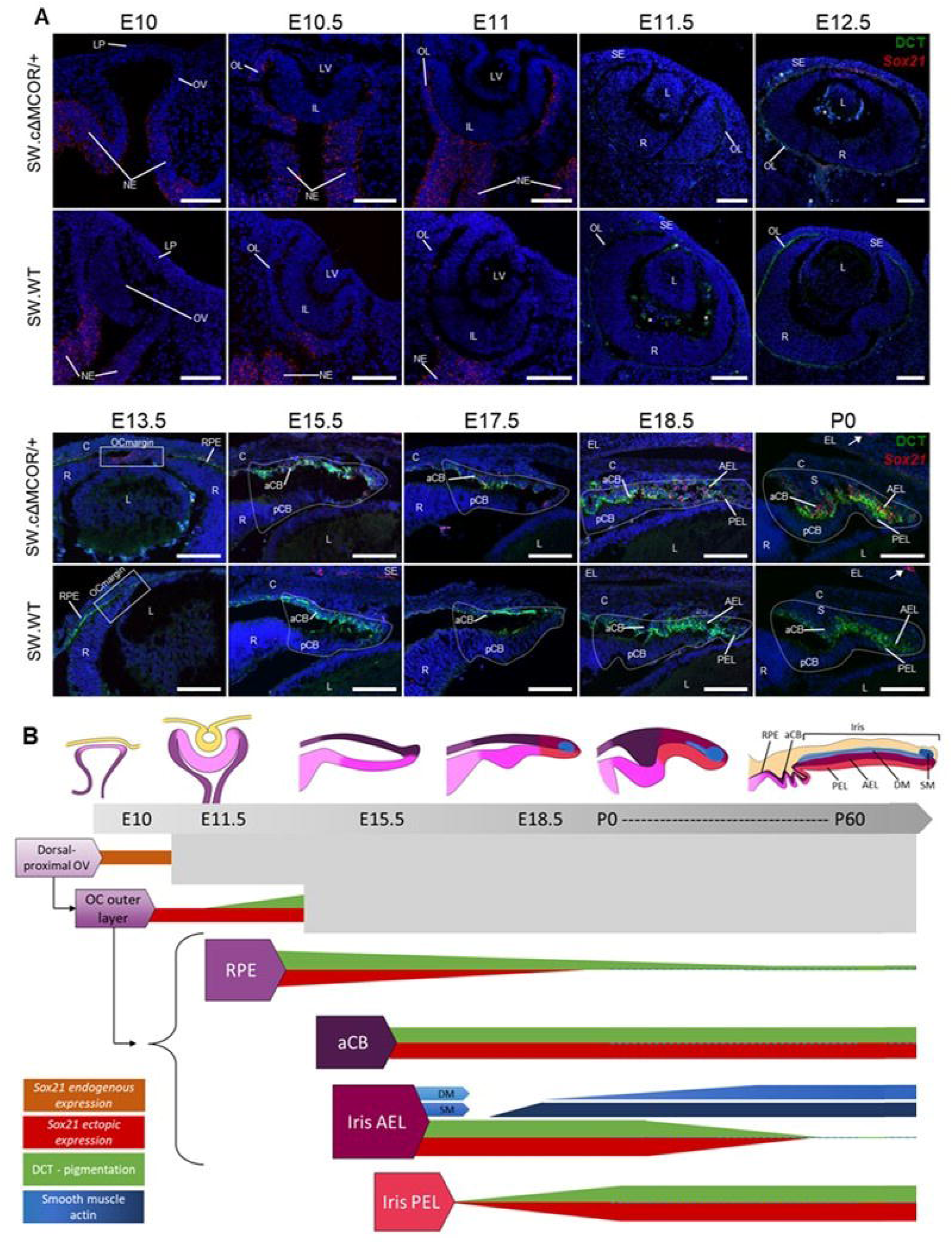
Expression pattern of *Sox21* and DCT during eye development. The spatio-temporal pattern of expression of *Sox21* and DCT was assessed using in situ hybridization (ISH) combined with immunofluorescence on multiple heads at various developmental stages, spanning from the invagination of the optic vesicle into the optic cup to the formation of the iris and CB (**A**). In the images, *Sox21* mRNA is stained in red, DCT in green, and autofluorescence, predominantly caused by blood vessels, is visible in cyan (white asterisks). The overall expression dynamics of *Sox21* and DCT during iris development are summarized in the scheme below (**B**). *Sox21* is expressed in the proximal region of the developing optic vesicle (OV) (depicted by the orange bar) at embryonic day 10 (E10) and is absent during the subsequent stages of optic cup (OC) formation. Around E15.5, endogenous *Sox21* is also observed in the eyelid and hair follicles (white arrows). In SW.cΔMCOR/+ animals, *Sox21* ectopic expression emerges from E10.5, encompassing the entire outer layer of the OC in red. This layer gives rise to the retinal pigment epithelium (RPE), and the anterior epithelium of both the CB (aCB) and iris (AEL) at its margin (outlined by white dotted lines), around E15.5. *Sox21* aberrant expression precedes the appearance of DCT (green) in the developing eye, which might be explained by enhancer priming preceding the initiation of gene transcription.^32–34^ As the RPE differentiates, the reduction in DCT is accompanied by a decline in *Sox21* during the late stages of development. The expression of both genes remains stable in the aCB from its development to maturation. Concurrently, the evolution in iris pigmentation aligns with the change in *Sox21* expression: the iris AEL is initially heavily pigmented (rich in DCT) and loses its melanin, along with *Sox21*, during postnatal maturation. The iris PEL progressively gains melanin in a centrifugal manner during development, becoming fully pigmented in adulthood, while also expressing *Sox21*. Notably, the sphincter and dilator muscles (depicted in blue) begin forming around E17.5 and E18.5, respectively, with the dilator maturing long after birth. The shaded box indicates the period from birth to the age of 2 months, during which we currently lack information regarding iris AEL pigmentation and the precise stages of dilator muscle (DM) development. **aCB:** ciliary body anterior epithelium, **AEL:** iris anterior epithelium, **C:** cornea, **DM:** dilator muscle, **EL:** eyelid, **IL:** optic cup inner layer, **L:** lens, **LP:** lens placode, **LV:** lens vesicle, **NE:** neuroectoderm, **OC:** optic cup, **OL:** optic cup outer layer, **OV:** optic vesicle, **pCB:** ciliary body posterior epithelium, **PEL:** iris posterior epithelium layer, **R:** retina, **RPE:** retinal pigment epithelium, **S:** iris stroma, **SE:** surface ectoderm, **SM:** sphincter muscle. Scale bars: **100µm**.

The absence of *Sox21* expression in the non-pigmented posterior epithelium layer (PEL) of both the developing and adult CB, along with the loss of both *Sox21* and DCT expression in the AEL of adult irises compared to developing irises (**Figure 3D and Figure 4**), further strengthens the evidence for the co-expression of *Sox21* and DCT in SW.cΔ MCOR/+ eyes (**Figure 4A and B**).

### The integrated analysis of SOX21 targets and gene expression deregulation in the MCOR-mouse indicates disruptions in genes related to iris development and symptoms associated with MCOR

We investigated the impact of the loss of the critical MCOR-causing region on the expression of genes lying outside of TADs 1 and 2 by analyzing RNAseq data from the new-born B6.cΔ MCOR/+ and B6.WT. This study detected 2500 differentially expressed genes (DEG ≥ 1.5 fold change (FC), p<0.05; **Figure 5A**). Gene Ontology (GO) enrichment analysis revealed a significant enrichment of genes associated with crucial processes such as cellular commitment, development, differentiation, and migration, particularly in the context of neural and sensory systems. Notably, the top pathways identified include cell fate commitment, forebrain development, regulation of cell differentiation, retina morphogenesis, regulation of neuron differentiation, and neuron projection development (**Figure 5B**). Consistent with these findings, previous research has highlighted the role of SOX21 in regulating adult neurogenesis *in vivo*, contributing to the generation of new neurons by repressing *Hes5* expression.^17^ Notably, our study reveals a downregulation of *Hes5* in the B6.cΔ MCOR/+ irises (FC = 0.56, p < 0.01), aligning with the established regulatory function of SOX21. Our study revealed additional neurogenesis-associated genes with altered expression and direct interaction with the *Sox21* product (**Figure S3**), notably SOX2 (MIM* 184429) which plays a crucial role in eye development, specifically anterior segment formation.^2021–25^ (FC = –1.85, p < 0.01; **Figure S3**)

**Figure 5.**
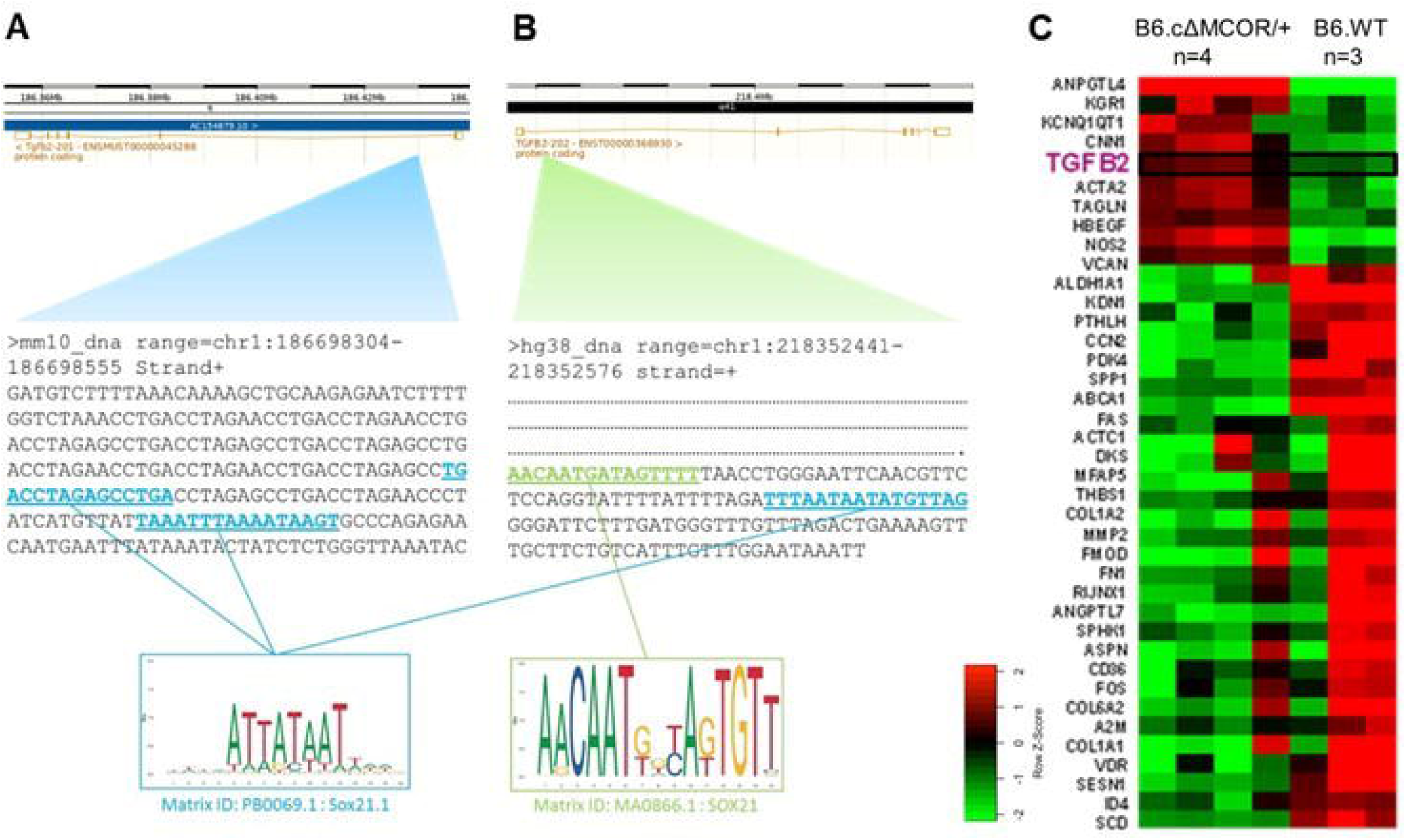
SOX21 consensus binding sequences in mouse and human *TGFB2* gene and RNA-seq analysis of the *Tgfb2* signaling pathway in the B6.cDMCOR/+ mouse as compared to B6.WT counterparts. A) ChIP-seq experiment reveals the DNA sequence binding to SOX21 in the mouse, particularly in the first intron of the *Tgfb2* gene. **B**) The syntenic sequence from the human genome assembly GRCh38, situated in the first intron of the *TGFB2* gene, is presented. The SOX21-consensus motif, according to the JASPAR database, is displayed at the bottom of the figure. The mouse binding sequence (PB006.1) of SOX21 is highlighted in blue, found in both the human and mouse sequences in the intron. The human motif (MA0866.1), shown in green, is also present in the *TGFB2* intron 1, suggesting a potential binding of SOX21 in this location. **C**) The RNA-seq heatmap shows genes of the TGFB pathway. While some variations are observable among individuals within the same group*, Tgfb2* is consistently overexpressed in all four cΔ MCOR/+ irises.

Furthermore, we noted the upregulation of key genes linked to major signaling pathways in iris development, including Wnt/β –catenin, TGFB, and Bone Morphogenetic Proteins (BMPs) signaling pathways (**Figure S4 and S4C**).^1^ Our investigation into the Wnt/β –catenin pathway highlights the substantial elevation of *Wnt2b* expression (MIM*601968; FC= 3.57, p < 0.01 **Figure S4A and S4C)**, a gene crucial for specifying iris progenitor cells towards a myoepithelial fate.^1^ In the context of BMP and TGFB signaling pathways, we observed upregulation in *Bmp7* (MIM112267; 1.92, p<0.05 **Figure S4B and S4C**), implicated in iris smooth muscle generation,^1^ and *Tgfβ2* (MIM*190220; FC 1.6, p<0.01), known for its role in iris and ciliary body specification, as well as in glaucoma^9–12^ (**Figure S4B and S4C**), and potentially associated with high myopia.^13–14^

Furthermore, we observed an enrichment of genes associated with the regulation of smooth muscle cell proliferation, though with a lower significance ranking (top 45; **Figure 5**). Notably, our analysis unveiled a decrease in expression of the *Des* gene (FC –2.0, p = 0.012), encoding desmin (MIM*125660) intermediate filaments known to be deficient in the iris anterior epithelium layer of MCOR-affected individuals.^3,26^ To explore if the smaller pupil size in B6.cΔMCOR/+ is linked to reduced DES levels, we used immunohistochemistry with DES and smooth muscle actin (SMA) antibodies as control on mouse iris sections. SMA stained both sphincter and dilator muscle fibers (**Figure S2**), but DES antibodies failed to detect the protein in both B6.cΔMCOR/+ and B6.WT irises.

### *Tgf****β****2* is a direct target of SOX21 in the iris of the MCOR-mouse model

To search for SOX21 target genes among the deregulated ones, we conducted ChIP-seq analysis using the SOX21 antibody on irises from newborn B6.cΔ MCOR/+ mice. This analysis uncovered 25 DNA regions throughout the genome (p ≤ 0.1), comprising 10 intragenic (all intronic), 14 intergenic, and 1 in promoter-TSS regions, respectively (Table 1). We examined deregulated expression (≥ 1.5 fold; p ≤ 0.05 B6.cΔ MCOR/+ *versus* B6.WT animals) among these genes and the nearest genes in cases where the binding occurred outside a gene. This search identified a unique gene: *Tgfβ2*; the remaining genes either exhibited no expression in the iris or showed no deregulation (**Table 1**). SOX21 binding to *Tgfβ2*, as determined by ChIPseq, was unequivocal (p<0.0001) and strongly supported by JASPAR analysis, which identified a consensus SOX21-binding sequence in the 252 bp intronic region specified by ChIPseq (GRCm38/mm10_chr1:186,698,304 – 186,698,555; 5.9 Kb downstream from the consensus donor splice-site of the 16 kb-long intron 1; **Figure 5A**). Notably, this sequence is conserved in the human *TGFΒ2* intron 1 orthologous region, containing numerous potential transcription factor-binding sites (GRCh38/hg38_chr1:218,352,441 – 218,352,576; **Figure 5B**). In line with the association between SOX21 and *Tgfβ2*, RNAseq analysis revealed dysregulation of the TGFB2 signaling pathway in the iris of B6.cΔ MCOR/+ compared to B6.WT littermates (**Figure 5C**).

**Table 1.**
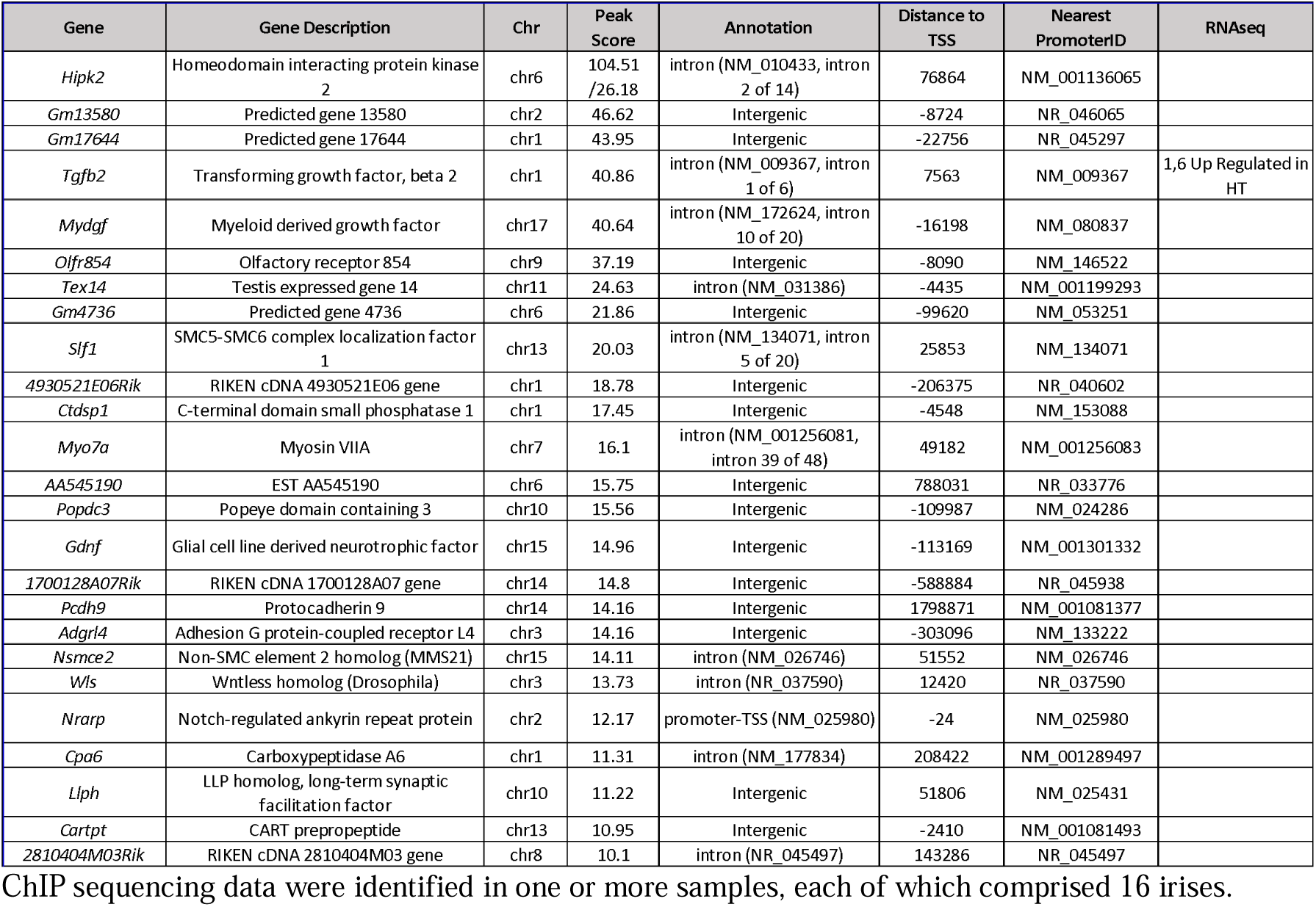
DNA regions identified by ChIPseq analysis for SOX21 binding in the iris of B6.c. Δ**MCOR/+ mice.** The genes listed in this table are restricted to those associated with a Peak score greater than 10, corresponding to a p-value of 0.1 or higher. The peak location is presented relative to the genomic distance from the transcription start site (TSS). Specifically, the 25 sites from the SOX21

### TGFB2 concentration is elevated in the AH of a MCOR subject

The ectopic expression of *Sox21* in the CB, along with SOX21 binding to a regulatory region of the *Tgfβ2* and its upregulation in B6.cΔ MCOR/+ irises, prompted us to measure TGFB2 concentration in the AH of 12-month-old B6.cΔ MCOR/+ and B6.WT mice using an ELISA assay. Concentrations, measured from pooled AH from both eyes, exhibited high variability in both WT and mutant eyes. Analysis of a large number of samples revealed no statistical difference between the two groups (**Figure S5**).

However, when analyzing AH samples collected on the same day during cataract surgery in a 45-year-old non-glaucomatous adult MCOR individual from a large French pedigree,^27^ as well as in eleven non-MCOR and non-glaucomatous individuals, we observed a significant increase in TGFB2 levels in the subject’s AH compared to the controls (**Figure 6**). Notably, the TGFB2 concentration was elevated as well in AH sample collected during cataract surgery in the fellow eye of the MCOR individual, which occurred two weeks after the first eye surgery. This observation provides additional support for the connection between SOX21 and TGFB2, emphasizing the role of TGFB2 in MCOR.

**Figure 6.**
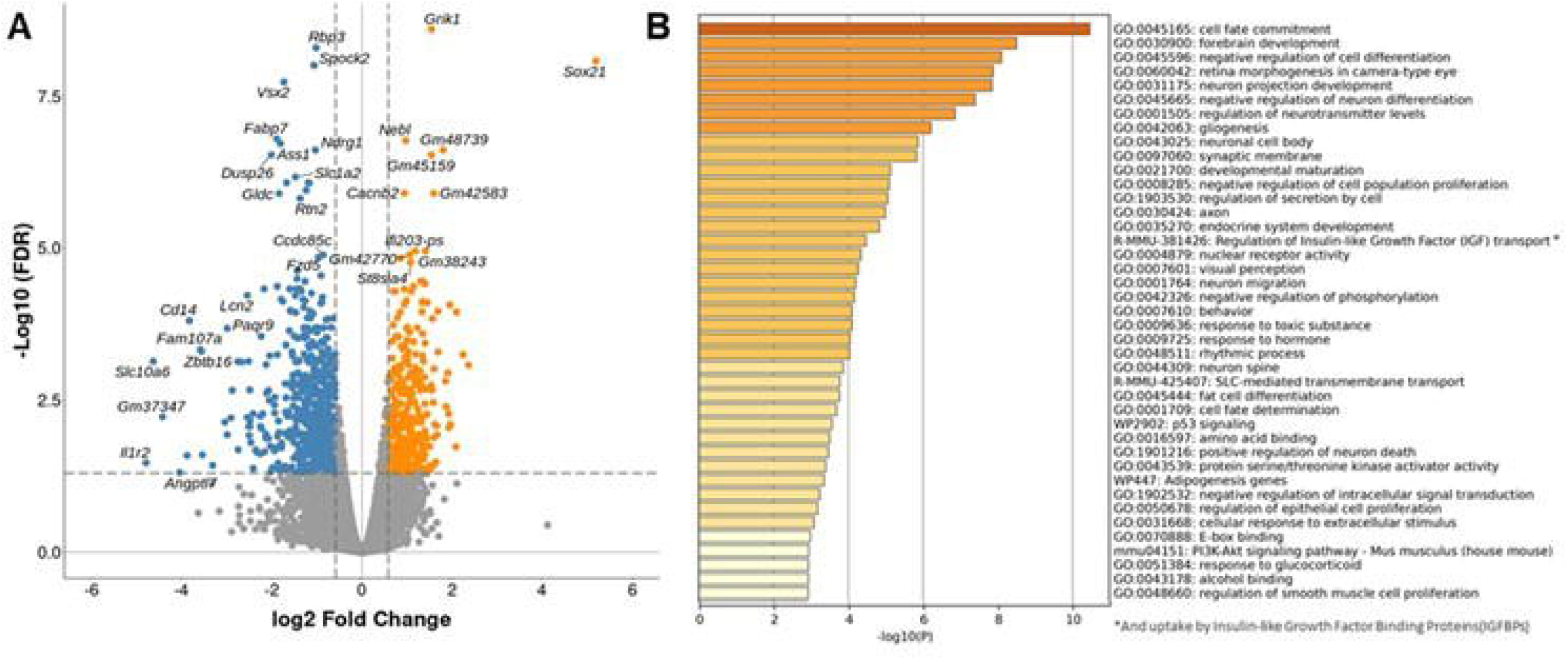
**Comparative analysis of gene expression in the iris of B6.cDMCOR/+ versus B6.WT littermates. A**) Volcano plots of the most differentially expressed genes (DEGs) from RNAseq data of B6.WT (*N*□ =□ 3) and B6.cΔ MCOR/+ (*N*□ =□ 4) irises at P0. On the x-axis (log2 scale), the difference in gene expression Fold Change (FC) is displayed and False Discovery Rate (FDR) adjusted significance is plotted on the y-axis (log10 scale). Negative values indicate down regulation; positive values indicate up regulation. **B**) Enrichment analysis in the category Biological Processes for most relevant DEGs using Metascape. The input DEGs lists are shown in a heatmap with enriched terms, which are colored by p-values.

**Figure 7.**
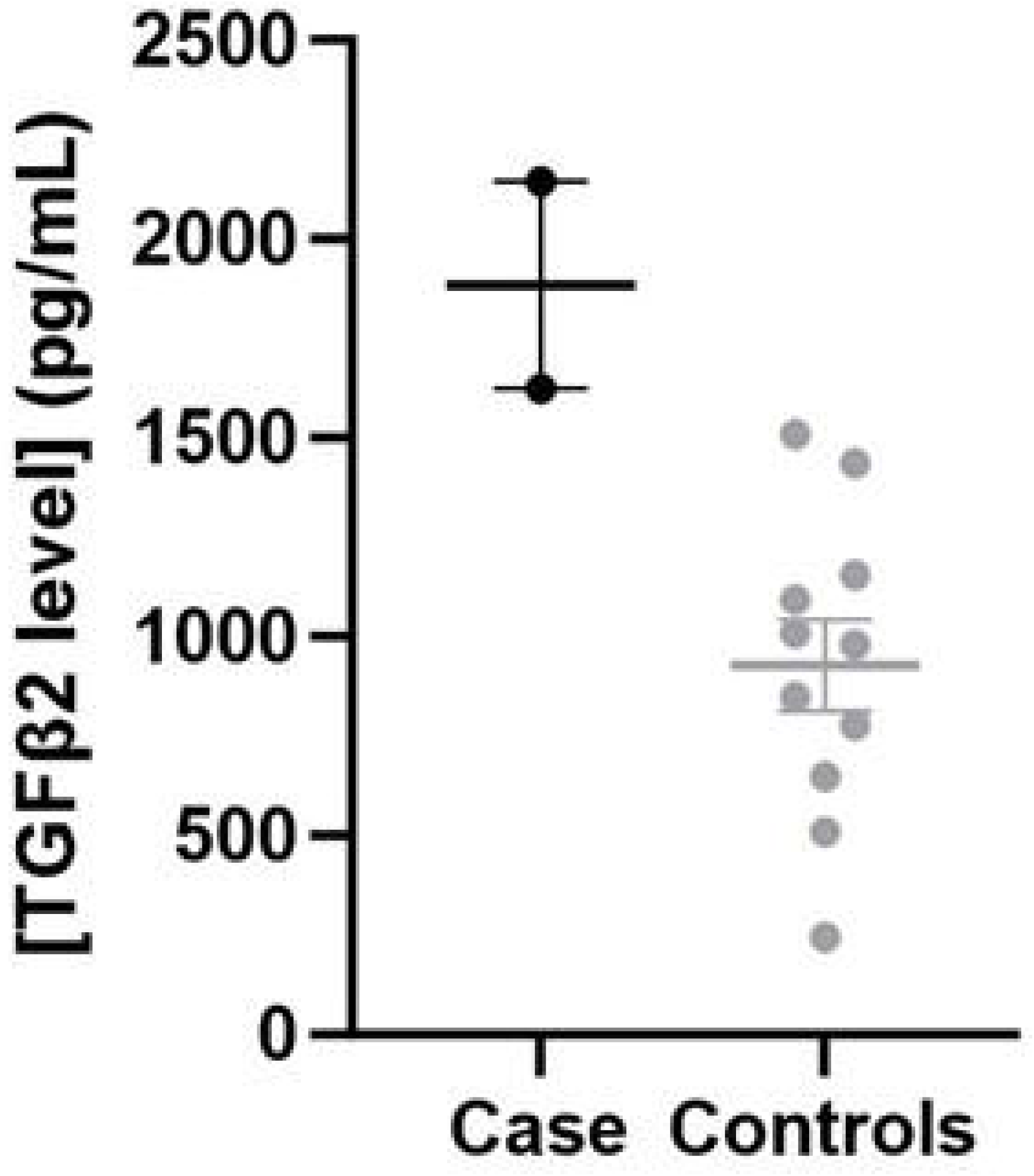
Analysis of TGFB2 concentration in the aqueous humor of a MCOR subject and controls. The ELISA dosage of TGFB2 in human samples reveals an accumulation of the protein in the MCOR subject analyzed, contrasting with 11 controls. Each data point on the graph represents the dosage value from the AH of an individual eye.

## DISCUSSION

Anatomical analyses of post-mortem eyes and iris specimens from MCOR-affected individuals show substantial iris thinning, stromal atrophy, and a lack of stromal contractile processes in the anterior iris epithelium, indicating a connection between iris closure and defective musculature development.^3^ Despite these findings, the underlying mechanisms and the association of the disease with abnormally long eyes and high IOP remain unclear. We leveraged the recent identification of the genetic etiology of the disease and the accessibility of CRISPR-Cas9 technology to develop a mouse model of the disease, analysis of which provides support for the central role of the SOX21 transcription factor in the pathology.

The role of SOX21 in eye development was documented in the chick^16^ and zebrafish^28^ but not in the mouse. In the chick, *Sox21* is transiently activated during the early stages of optic vesicle morphogenesis and specification in the lens and retina but is not expressed thereafter;^16^ *Sox21* expression in the eye ceases before the onset of iris development. Loss of *Sox21* function in the chick, similar to findings in zebrafish, disrupts normal lens development,^29^ whereas it has no effect on the mouse eye,^30^ which aligns with our immune histological analysis detecting no expression of SOX21 in the anterior segment of the developing mouse eye.

In our study, we show that deleting the critical region in the mouse genome, equivalent to MCOR in humans, results in constitutive *Sox21* expression. Remarkably, we observe colocalization of *Sox21* and the melanogenic marker DCT in both developing and adult eyes. According to the predicted 3D structural organization, *Dct* and *Sox21* are located in two adjacent neighboring TADs, the boundaries of which are disrupted by MCOR-causing deletions, while the genes themselves remain unaltered. In line with prior research associating TAD disruptions with developmental defects,^31^ we propose that the loss of TAD boundaries in MCOR may initiate a reorganization, potentially activating *Sox21* through *Dct* enhancers. The delayed detection of the DCT protein (E11.5) compared to the detection of *Sox21* mRNA (E10.5) is consistent with studies demonstrating that enhancer priming precedes the initiation of gene transcription.^32–34^

In line with the melanogenic DCT protein’s expression pattern, we detected ectopic expression of *Sox21* in the pigmented PEL of the iris in adult MCOR-mouse model, while it was absent in the AEL responsible for dilator muscle formation.^35–36^ Contrastingly, during embryonic development, the iris AEL is pigmented and expresses DCT. While literature lacks reports on iris muscle development in mice, in rats, the dilator muscle initiates development around E18.5 and continues postnatally.^35^ In our study, we found DCT in the iris AEL of E18.5 and P0 WT mice, and both DCT and Sox21 in MCOR-mice. This suggests that abnormal *Sox21* expression in the AEL of the MCOR-mouse iris coincides with the onset of dilator muscle development and decreases during postnatal development, along with the loss of DCT expression (**Figure 4**). These findings might elucidate histological observations of microcoric iris specimens, displaying poor differentiation, disordered fibers lacking myofilaments and intermediate filaments, or even complete growth inhibition, which suggest issues in the terminal differentiation stages of the anterior iris epithelium (reviewed by Angee et al.^3^). In the mouse, our histological analysis failed to detect such anomalies. This could be attributed to the minimal iris phenotype of the mutant mouse, potentially linked to weaker *Sox21* expression, divergent timelines in iris dilator muscle development, or dissimilarities in the 3D architecture of the region encompassing the MCOR locus between humans and mice. While the iris phenotype in the mutant mouse is minimal, possibly due to the dilator muscle playing a less significant role in mice compared to humans, it is noteworthy that we observed a reduced expression of the desmin gene in the iris of newborn mutant mice. While we couldn’t directly confirm the reduced abundance of the DES protein product in the iris AEL due to the lack of suitable antibodies for immunocytochemistry analysis, this observation aligns with previous studies on iris anatomy. These studies have reported the absence or severe reduction of desmin fibers in the anterior epithelium of individuals with microcoria.^3,26^ Furthermore, RNAseq analysis revealed the deregulation of key genes known to be crucial in signaling pathways important for iris development, such as *Wnt2b*. In the early stages of eye development in chicken and mouse embryos, *Wnt2b* expression is concentrated at the marginal tip of the developing retina, with a notable increase in the dorsal part of this region, responsible for forming the pigmented epithelium of the iris.^1^ Interestingly, this specific region coincides with the location of ectopic expression of *Sox21* observed in B6.cΔ MCOR/+ mice. (**Figure S4A**). The lack of SOX21 protein binding to *Wnt2b* regulatory sequences, as revealed by ChIPseq analysis, suggests that the deregulation is likely indirect to *Sox21* induction. Nonetheless, the observed spatial correlation between *Sox21* expression and the regional modulation of *Wnt2b* provides valuable insights into the regulatory mechanisms governing early eye development and the formation of the iris pigmented epithelium. Among the deregulated genes identified by RNAseq, *Tgfβ2* is the sole gene found to bind SOX21 in mutant irises. This binding occurs through a consensus binding sequence located in intron 1. This observation aligns with a growing body of evidence across various organisms, suggesting that for a subset of genes, the critical sequences regulating expression are not primarily situated in the promoter but within introns within the first kilobase of transcribed sequences.^37^

Presently, the specific mechanisms responsible for the observed default in muscle development, observed in both the mouse model and individuals affected by MCOR, remain elusive. One plausible hypothesis suggests the involvement of TGFB2. This factor is notably secreted within the periocular mesenchyme and plays a pivotal role in the development of various ocular components, including the anterior chamber –critical for AH outflow-and the stroma of the iris and CB.^38^ Although the iris dilator muscle originating from the optic cup margin, derived from neuroectoderm, its myofibers manifest within the stroma. Therefore, an aberrant expression of *TGFB2* within the iris could potentially disrupt stromal development, subsequently affecting the proper formation of the dilator muscle within this context. This proposed link underscores the importance of investigating the interplay between TGFB2 signalling and muscle development in the iris, offering a potential avenue for understanding the muscle-related anomalies observed in the context of MCOR. Remarkably, several studies reported significantly elevated levels of TGFB2 in the AH of individuals with POAG, as well as in cultured glaucomatous cell strains and isolated human glaucomatous trabecular meshwork (TM) tissues.^10,11^ The cause and cellular source of TGFB2 in glaucomatous eyes remain elusive. However, studies have demonstrated that cells from the TM, through which AH is drained out at the iridocorneal angle, express an active TGF receptor complex and respond to exogenous TGFB2.^39,40^ This has been reported to result in increased synthesis of extracellular matrix proteins (ECM), accumulation of which increases resistance to aqueous outflow, leading to elevated IOP and GLC in human, primates, cats and mouse.^9–12,41^

This knowledge, combined with (i) the histopathologic examination of TM biopsies from two microcoric brothers with elevated IOP, that revealed ECM accumulation,^42^ and (ii) an elevated concentration of TGFB2 in the AH of an individual with MCOR, implies that GLC associated with MCOR may be initiated by SOX21. The strength of this inference is heightened by our findings, which provide evidence of ectopic *Sox21* expression in the CB – where the AH is produced-specifically in the non-pigmented PEL.^43^ Furthermore, a recent study has uncovered a significant correlation between elevated TGFB2 levels in the AH and axial elongation.^13,14^. This discovery not only suggests a connection between TGFB2 and, by extension, SOX21, but also extends the potential link to high myopia.

Finally, we demonstrate that homozygosity for the critical MCOR-causing deletion is lethal in the mouse. While the possibility of *Sox21* playing a role cannot be ruled out, it is more likely that lethality results from the homozygous deletion of *Tgds*. Notably, biallelic mutations in *TGDS* have been documented to lead to a severe developmental disorder characterized by the Pierre Robin anomaly, known as Catel-Manzke syndrome (MIM# 616145). Intriguingly, to our knowledge, no instances of homozygosity or compound heterozygosity for loss-of-function alleles was reported, suggesting that such a situation could be lethal. An alternative explanation could consider the aberrant accumulation of pyknotic nuclei in developing ectodermal and neuroectodermal derivatives such as rhombomeres, neural crest-derived cranial ganglia, and placodes observed as a cause of lethality in mutant embryos. From a morphogenetic standpoint, the pathophysiological accumulation of pyknotic cells is linked to embryotoxicity,^15,44^ attesting of teratogenic or pathogenic conditions and accounts for cell degeneration initiated by apoptosis. In neuronal and glial-derived tissues in the central and peripheral nervous system, pyknosis is interpreted as signs of cell degeneration that could be enhanced through the activation of Ca++-activated proteases^45^ or metalloproteases.^46^ Notably, tissues exhibiting intense accumulation of pyknotic cells coincide with active production of TGFB2 during normal development.^47–49^ In mice lacking *Tgfβ2*, inhibiting this pathway prevents neuroblastic lineages from apoptosis.^50^

In conclusion, our study demonstrates that the critical submicroscopic deletion associated with MCOR potentially disrupts the 3D architecture of the region, leading to modified gene interactions. This alteration results in the ectopic expression of the Sox21 transcription factor in *Dct*-expressing pigmented cells within the iris and CB of MCOR mice. Importantly, we posit that in MCOR, both GLC and high myopia stem directly from SOX21-mediated TGFB2 overexpression in the CB, rather than arising due to abnormal iris development. Consequently, we propose that SOX21 serves as a crucial link, connecting iris malformation, high myopia, and GLC in MCOR. This positions MCOR as an invaluable model for scrutinizing eye development and unraveling the underlying mechanisms of common myopia and POAG.

## MATERIALS AND METHODS

### Generation of a MCOR-mouse model

The B6.cΔ MCOR strain utilized in this study was generated through CRISPR/Cas9 methodology at the Transgenesis Platform of the Animal Facility (LEAT) at the Imagine Institute (Paris). Animal procedures received approval from the French Ministry of Research and were conducted in compliance with the French Animal Care and Use Committee from the University of Paris Cité (APAFIS #14311-201801151627355). Two guide RNAs (sgRNAs, 5’-CTCACAGTTTGGTCCAGGCT-3’ and 5’-ATTCCCCAGCAGAGAGGCGC-3’) targeting the MCOR locus were designed using the CRISPOR web tool (http://crispor.tefor.net/). Deleterious alleles were induced via CRISPR/Cas9 ribonucleoprotein (RNP) complex microinjection into C57BL/6 mouse zygote pronuclei, as described previously.^51^ Offspring were genotyped from genomic DNA through PCR amplification using appropriate primers (forward 5’-GGATGTGCGTTCAAATCCCAGTACC-3’ and reverse 5’-TGTTCCCATGCAGTGTGGCAATG-3’). To eliminate potential off-target mutations, transgenic animals underwent multiple generations of backcrossing with C57BL/6 mice. SW.cΔ MCOR/+ animals were obtained by crossing B6.cΔ MCOR/+ with Swiss albino mice over several generations.

### Pupillometry analysis

Two-month-old mice (8 B6.WT and 8 B6.cΔ MCOR/+) underwent dark adaptation overnight, and pupillary responses were recorded following established protocols (Kostic et al. 2016). To avoid the potential effects of anesthesia, animals were not anesthetized. The baseline pupil diameter was defined as the mean pupil diameter during the 500 ms before light onset. Subsequently, all pupil sizes were normalized relative to this baseline value. The light stimulus sequence comprised 50 ms exposures to (0.01, 0.1, and 3.16) W/m2 white light and a 20s exposure to 1 W/m2 blue light. Pupil diameter was automatically determined using Neuroptics A2000, Inc. software. One-way/two-way ANOVA analyses were employed to identify significant differences.

### RNA extraction

Mice aged from 0 days postnatal (P0) to 12 months, were euthanized, and their eyes were enucleated. On ice, the eyes were dissected to collect the iris and CB. Each eye was halved following the ora serrata, and after lens removal, the iris and CB were extracted from the anterior segment. Tissues from both eyes were pooled for total RNA extraction using the RNAeasy MiniKit (Qiagen, Hilden, Germany), following the manufacturer’s instructions, post-mechanical tissue disruption. RNA quality and concentration were assessed through optical density measurements (Nanodrop, Thermo Fisher Scientific, Waltham, MA, USA), and RNA integrity was analyzed using capillary electrophoresis with a Tape Station (Agilent, Santa Clara, CA, USA) and High Sensitivity RNA Screen Tape.

### RNAseq analysis

The Ovation Mouse RNA-Seq System from NuGEN (Tecan Genomics, Leek, NL) was employed to prepare RNA-seq libraries using 25 ng of total RNA, following the manufacturer’s recommendations. This kit enables strand-specific RNA-Seq library construction using 10 to 100 ng of total RNA and utilizes the Insert Dependent Adaptor Cleavage (InDA-C) technology to eliminate ribosomal RNA transcripts. An equimolar pool of the final indexed RNA-Seq libraries was sequenced on an Illumina HiSeq 2500, targeting a sequencing depth of approximately 30 million paired-end reads per library. On average, 28 million reads were obtained. FASTQ files were mapped to the ENSEMBL Mouse (MM10) reference using Hisat2 and counted by feature. Counts from the Subread R package. Read count normalizations and group comparisons were conducted using three independent and complementary statistical methods (Deseq2, edgeR, LimmaVoom). Flags were generated from counts normalized to the mean coverage, where counts <20 were considered as background (flag 0) and counts >=20 were considered as signal (flag=1). P50 lists, utilized for statistical analysis, grouped genes showing flag=1 in at least half of the compared samples. Results from the three methods were filtered at p-value <= 0.05 and fold changes of 1.2/1.5/2, and grouped by Venn diagram. Cluster analysis employed hierarchical clustering with the Spearman correlation similarity measure and ward linkage algorithm. Heat maps were generated using the R package ctc: Cluster and Tree Conversion and visualized with Java Treeview software. Functional analyses were conducted using Ingenuity Pathway Analysis (IPA, Qiagen).

### RT-qPCR analysis

cDNA was synthesized using the cDNA Verso reverse transcription kit (Thermo Fisher Scientific). Quantitative RT-PCR (RT-qPCR) was conducted using SYBR Green (Sso Advanced Universal SybrGreen Supermix, Bio-Rad, Hercules, CA, USA), with *B2m*, *Tbp*, *Gusb* and *Hprt* as reference genes. The primer sequences for qPCR were as follows: *Sox21* forward 5’-AGCCTGTGGACCACGTCAA-3’ and reverse 5’-CCGACTCGGTGAGCAGCTT; *Dct* forward 5’-AGATTGTGTGCGACAGCTTG-3’ and reverse 5’-AAGGGAGGGCTGTCAAACTT-3’; *B2m* forward 5’-CCTGTATGCTATCCAGAAAACCCCT-3’ and reverse 5’-CGTAGCAGTTCAGTATGTTCGGCTT-3’; *Tbp* forward 5’-TGACCTAAAGACCATTGCACTTCGT-3’ and reverse 5’-CTGCAGCAAATCGCTTGGGA-3’; *Gusb* forward 5’-CTGGGGTTGTGATGTGGTCTGT 3’ and reverse 5’-TGTGGGTGATCAGCGTCTTAAAGT-3’; *Hprt* forward 5’-GTTGGTACAGGCCAGACTTTGTT-3’ and reverse 5’-AAACGTGATTCAAATCCCTGAAGTA-3’. The relative expression of *Sox21* and Dct was calculated using the method described previosuly^53^.

### Western blot analysis

The western blots were performed using 40 μ g of total iris protein lysates. The following antibodies were used in this study: SOX21 (AF3538, R&D Systems, Minneapolis, MN, USA) and beta-Actin (ab8227, Abcam, Cambridge, UK). Secondary antibodies, goat IgG or mouse IgG HRP (Thermo Fisher Scientific) were used at a 1:10000 dilution. Western blot membranes were developed using the Clarity Western ECL substrate (Bio-Rad), and the signal was detected with a Chemidoc MP Imaging System (Bio-Rad).

### ChIPseq analysis

The irises of 24 newborns (P0), including both B6.cΔ MCOR/+ and B6.WT mice, were collected for ChIP-seq analysis. Three samples were prepared for B6.cΔ MCOR/+ mice and three for B6.WT mice, with each sample comprising 16 irises. The ChIP assay was performed on all samples using 2 μ g of chromatin.

Cells were crosslinked with 1% formaldehyde for 10 minutes at room temperature. ChIP was carried out using the iDeal ChIP-seq kit for Transcription Factors (C01010055, Diagenode, Liege, Belgium), following the provided protocol. Immunoprecipitation utilized an anti-SOX21 antibody (AF3538, R&D Systems). Sequencing was performed on an Illumina HiSeq 4000, running HiSeq Control Software HD version 3.4.0.38. Quality control of sequencing reads was conducted using FastQC. Reads were aligned to the reference genome (GRCm38/mm10) from the UCSC genome browser using BWA software v.0.7.5a. All samples were filtered using a list of genomic regions created from the two input samples. Subsequently, samples were deduplicated using SAMtools version 1.3.1. Alignment coordinates were converted to BED format using BEDTools v.2.17, and peak calling was performed using MACS2.

### ELISA assay

Mice were euthanized at 5, 6, and 12 months old, and AH was collected from both eyes using a 27G-syringe inserted at the ora serrata (around 5 µL per eye). The AH from both eyes was pooled, and TGFB2 proteins from the samples were activated using one volume of HCl and two volumes of NaOH, as per the manufacturer’s instructions (Mouse TGF-beta 2 Quantikine ELISA Kit, R&D Systems). The samples were diluted (1:80), and the quantity of total inactive TGFB2 protein was measured using the Quantikine kit.

Human AH was collected from ten subjects undergoing cataract surgery at Clinique Jules Verne, Nantes, France. The mean age of the nine controls was 76.7 ± 5 years old, while the MCOR individual was the youngest at 44 years old. All samples were collected with the written and informed consent of the subjects. TGFB2 dosage in each eye (13 controls + 2 cases) was conducted with the Human TGF-beta 2 Quantikine ELISA kit (R&D Systems), following the provided instructions.

### Immunofluorescence and RNAscope analysis

Pregnant mice were euthanized to collect embryos at various stages, from E10.5 to P0. Accurate embryonic stages were assigned post-cutting by consulting established atlases due to variations in early embryo development within littermates. Embryo heads were fixed with 4% Paraformaldehyde (PFA), and fixed heads as well as fresh adult eyes were embedded in Optimal Cutting Temperature (OCT) compound and frozen in liquid nitrogen for rapid tissue freeze. OCT blocks were stored at –80°C and brought up to –20°C prior to cutting (CM3050S, Leica, Wetzlar, Germany). Frozen tissue sections (14µm thickness) were collected at the level of the pupillary aperture, air-dried, and stored at –80°C.

Frozen sections were rinsed in PBS to remove surrounding OCT, and antigen retrieval was performed using Citrate Buffer (pH 6.0) in a 95°C water bath for 30 minutes. Slides were then cooled for 30 minutes at room temperature. Tissues were blocked for 1 hour in PBS with 1% bovine serum albumin (BSA), labeled with primary antibodies anti-SOX21 (AF3538, R&D Systems), anti-DCT (ab74073, Abcam), and Cy3-coupled anti-alpha Smooth Muscle Actin (C6198, Sigma-Aldrich, Saint-Louis, MO, USA), followed by appropriate secondary antibodies. Slides were counterstained with DAPI and mounted with Fluoromount medium (Sigma-Aldrich).

For mRNA detection, slides were processed using the RNAscope Multiplex Fluorescent v2 Assay, and subsequent immunohistochemistry was performed. All images were acquired with a Spinning Disk Confocal microscope (Carl Zeiss, Oberkochen, Germany), capturing whole sections with a 10X objective and specific areas of interest (iris, CB, and margins of the optic cup) with a 40X oil-objective. Raw images were converted to TIFF for further analysis using Fiji (v1.53t).

### Statistical analysis

Unpaired bilateral Student’s t-test was used for two-group analyses. Two-way ANOVA analysis was used for multigroup analyses (GraphPad Prism, ver. 10.0.3). Data are presented as the mean ± SEM; p<0.05 was considered significant.

## ACKNOWLEDGEMENTS

This research has been generously supported by grants from the Agence Nationale de la Recherche (ANR# –20-CE 12-0019-01), the Institut Nationale de la Santé et de la Recherche Médicale (INSERM), MSD Avenir (DEVO-DECODE program), the Fondation Vision, and the Association Retina France.

## AUTHOR CONTRIBUTIONS

JMR and LFT designed the project. BP, JLV and PD generated the MCOR mice. CA and EE performed and interpreted the molecular, histological and imaging experiments. DH analyzed the Hi-Cseq data. CK and SVC performed the pupillometry experiments. ED, JK, PC, JP and NCh provided clinical data. MZ and CBF performed sequencing of the RNA samples. NCa performed bioinformatic analyses. BN and JC provided technical assistance. CA wrote the manuscript and prepared the figures with contributions from all coauthors. JM and LFT reviewed all of the data and edited the manuscript. ChatGPT has been used to edit English.

