## Supplementary data for "Modeling the critical MCOR-causing deletion in mouse unveils aberrant *Sox21* expression in developing and adult iris and ciliary body, and implicates *Tgfβ2* in MCOR-associated glaucoma and myopia"

**SUPPLEMENTARY FIGURES**

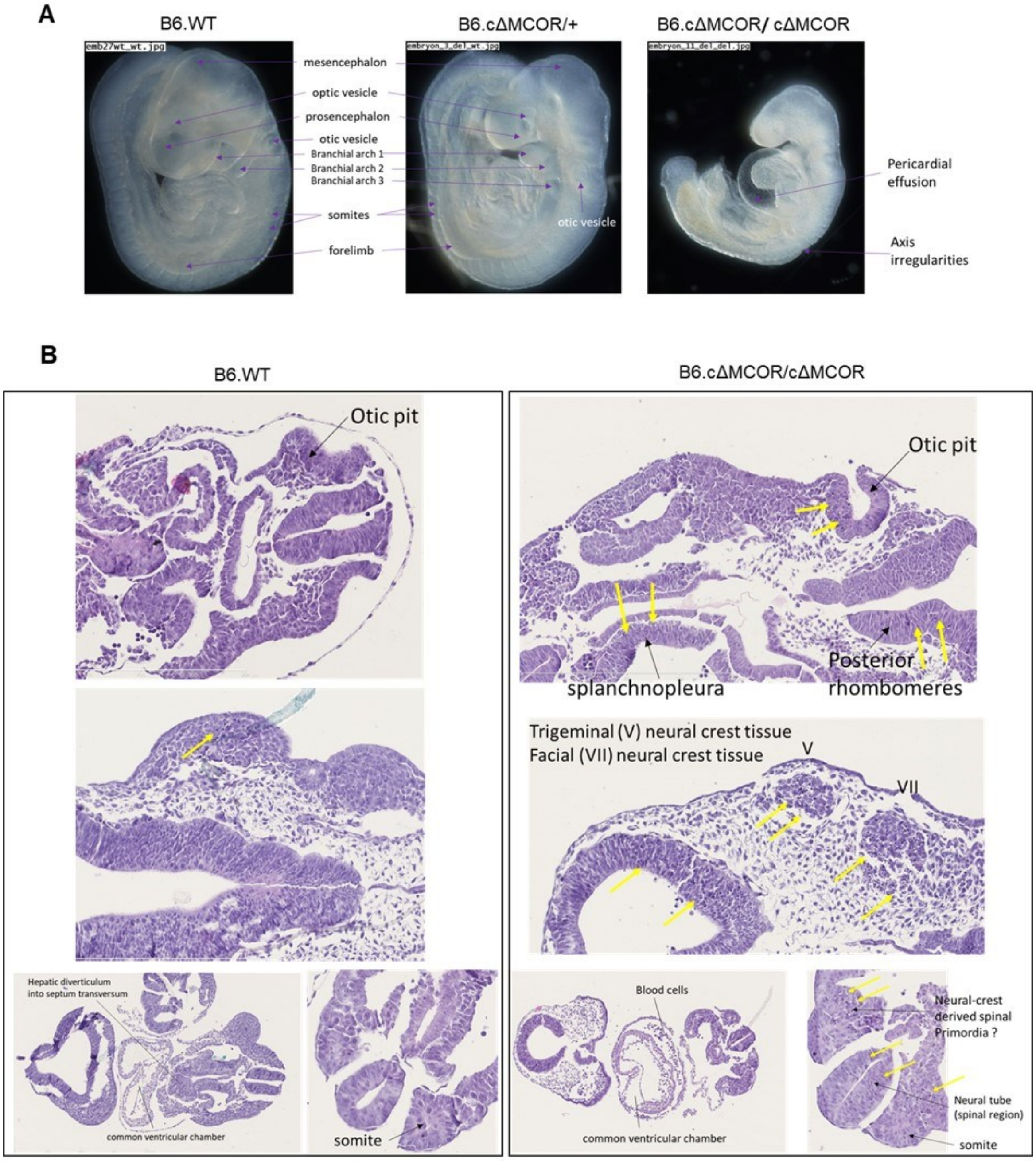


**Figure S1. Lethality studies. A**) Magnified pictures of embryos collected at 9.5 embryonic days. Genotyping was performed using the placenta. All individuals carrying the deletion in homozygosity presented a similar phenotype, with a smaller size and varying degrees of cardiac hypertrophy. Heterozygous and WT embryos presented no developmental defects. **B**) Histological analysis of homozygous embryos (E9.5) and comparison with WT counterparts. Mutant embryos exhibit an abnormal amount of pyknotic nuclei (yellow arrows) in numerous regions. Some pyknotic nuclei were found scattered in the neural tube and others were focalized as packets in particular regions such as for example posterior rhombomeres of the hindbrain, spinal region of the neural tube, and region of the optic vesicle. In the control embryo, some pyknotic nuclei can be observed in the neural tube but never at this amount. Packets of pyknotic nuclei were detected in the mesenchyme adjacent to the neural tube-containing pyknosis and in the somites. Pyknosis was also observed in the splanchnopleura tissue.

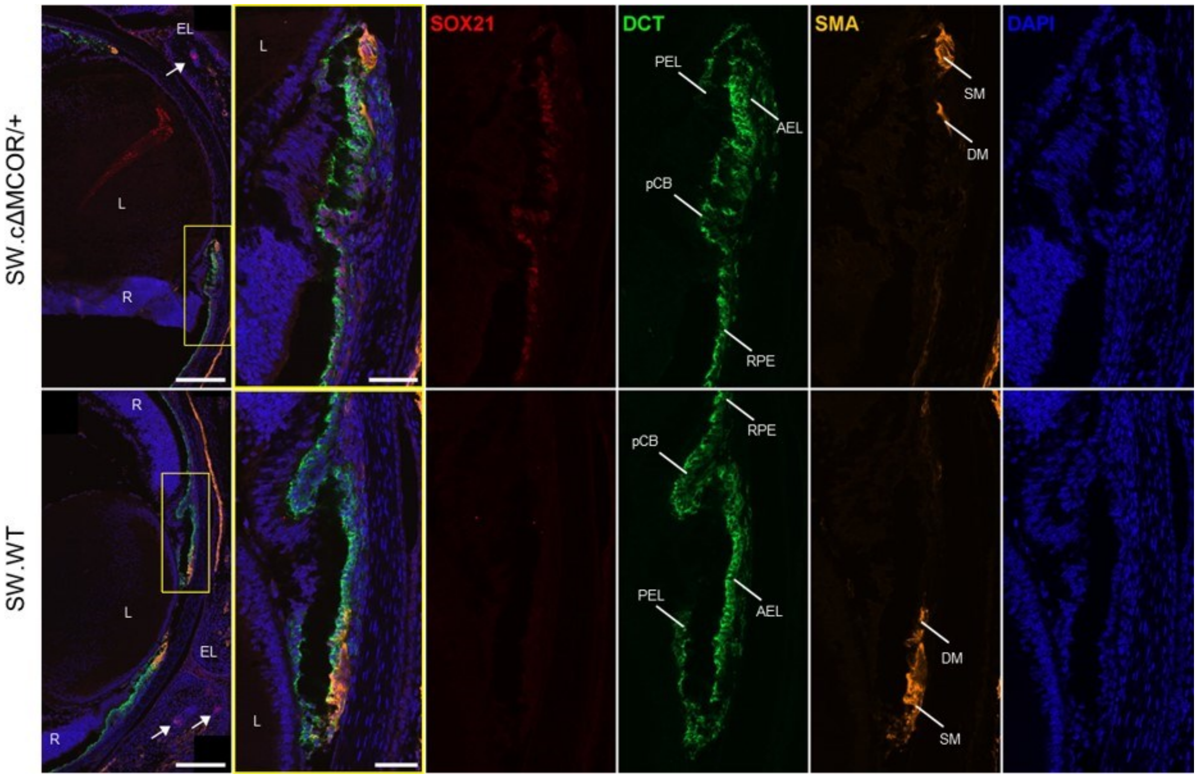


**Figure S2. Expression of SOX21, DCT and SMA at E18.5.** Global view of the anterior segment of the eye at late embryonic stage E18.5 (left) reveals the endogenous expression of *Sox21* in the eyelid and in hair follicles (white arrows). The yellow frame is represented with higher magnification on the right. In SW.cΔMCOR/+ eye, colocalization of SOX21 and DCT appears clearly in pigmented epithelia of the eye, namely the RPE, aCB, iris AEL and PEL. The SMA staining show the apparition of the first DM fibers behind the SM that developed earlier. No difference in both muscles were observed between SW.cΔMCOR/+ and WT counterparts. **AEL:** iris anterior epithelium layer, **DM:** dilator muscle, **EL:** eyelid, **L:** lens, **pCB:** posterior ciliary body epithelium, **PEL:** iris posterior epithelium layer, **R:** retina, **RPE:** retinal pigment epithelium, **S:** iris stroma, **SM:** sphincter muscle. Scale bars: **200µm** (left) and **50µm** (right zoom).

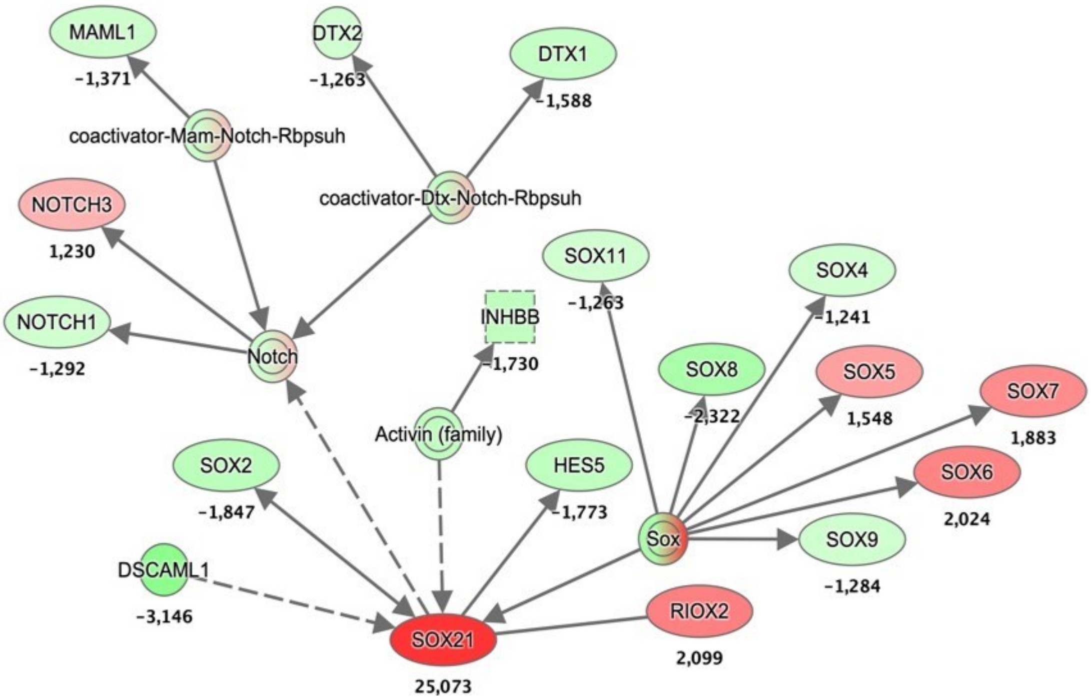


**Figure S3: SOX21 networks identified using differentially expressed gene analysis in cΔMCOR Mice.** Ingenuity Pathway Analysis (IPA) tools was used to construct a gene interaction network between SOX21 and other genes. The values next to the gene names refer to fold change of gene expression. Red nodes represent upregulated genes, while the green nodes are for downregulated genes. Edges (lines and arrows between nodes) represent direct (solid lines) and indirect (dashed lines) interactions between molecules as supported by information in the Ingenuity knowledge base.

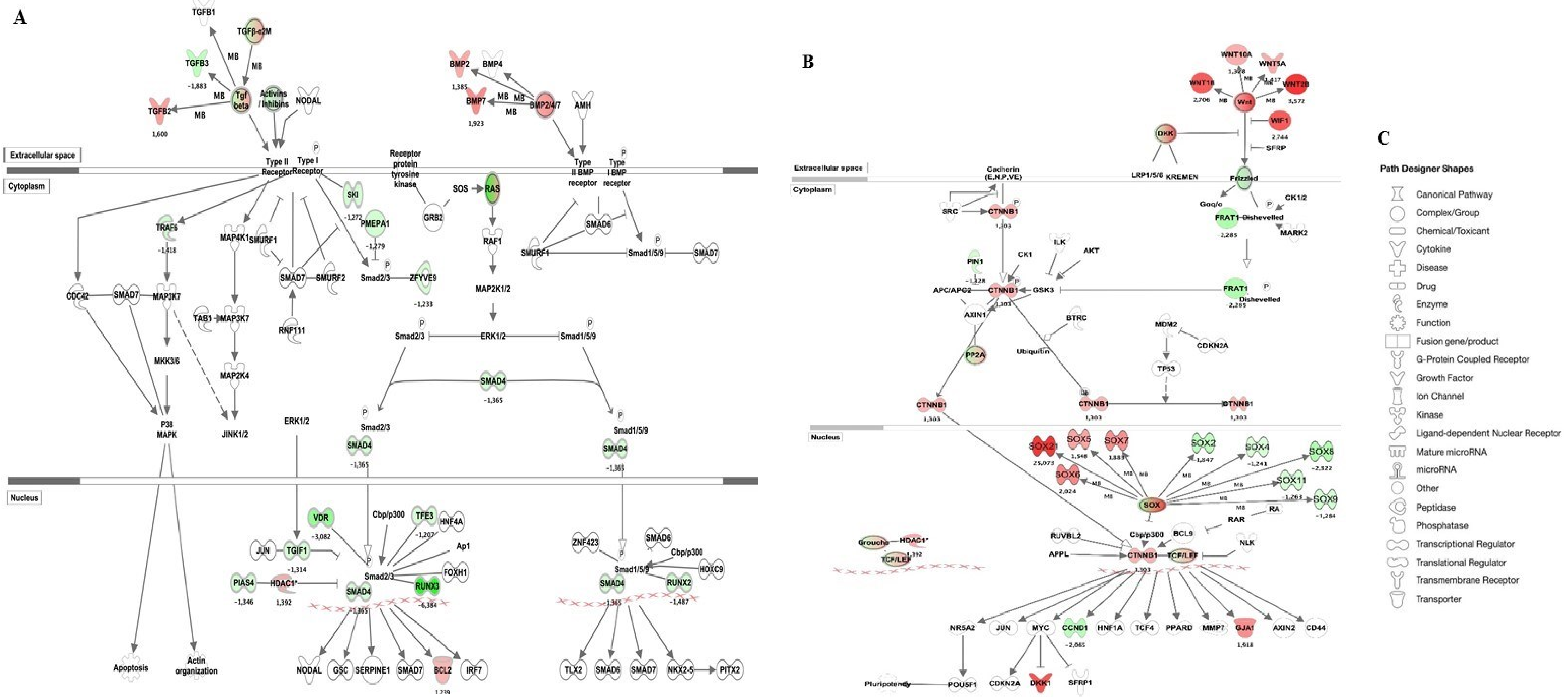


**Figure S4: Three canonical pathways selected by their potential roles in the physiopathology of the disease.** A) Wnt/β-catenin signaling pathways and B) Combined analysis of Transforming Growth Factor Beta (TGF-β) and Bone Morphogenetic Proteins (BMPs) signaling pathways. The results were filtered at a *p*-value ≤0.05 and a fold-change of 1.2. Gene lists analysed with three methods (Deseq2, Voom and edgeR) were uploaded to IPA to determine canonical pathways differentially regulated in the cΔMCOR irises compared to the B6.WT counterpart. Genes that are significantly up- and down-regulated are shown in red and green respectively. The intensity of the color corresponds to an increase or decrease in fold change levels of gene expression. Genes in white did not exhibit significant changes in expression. C) The pathways designer shapes of molecules is defined in the graphical legends.

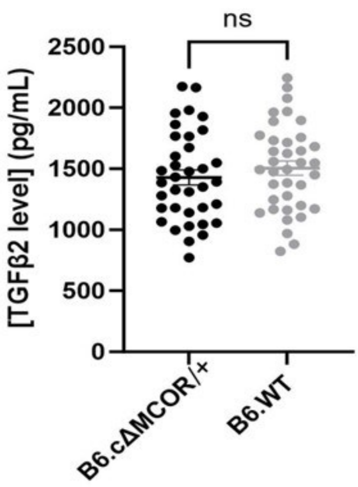


**Figure S5. Analysis of TGFB2 concentration in the mouse aqueous humor.** The ELISA dosage of TGFB2 in the aqueous humor (AH) of one-year-old mice utilized twenty animals for each genotype, with each dot on the graph representing an individual (AH from both eyes pooled due to limited sample quantity). The dosage results exhibit considerable variability in the measured TGFB2 levels in mouse AH. Notably, no significant (ns) difference was observed between the two groups.
